# An atypical role for the myeloid receptor Mincle in CNS injury

**DOI:** 10.1101/044602

**Authors:** Thiruma V. Arumugam, Silvia Manzanero, Milena Furtado, Patrick J. Biggins, Yu-Hsuan Hsieh, Mathias Gelderblom, Kelli PA MacDonald, Ekaterina Salimova, Yu-I Li, Othmar Korn, Deborah Dewar, I Mhairi Macrae, Robert B. Ashman, Sung-Chun Tang, Nadia A. Rosenthal, Marc J. Ruitenberg, Tim Magnus, Christine A. Wells

**Affiliations:** Department of Physiology, Yong Loo Lin School of Medicine, National University of Singapore, Singapore 117597, Singapore; School of Biomedical Sciences, The University of Queensland, Brisbane 4072, Australia; Australian Institute for Bioengineenng and Nanotechnology, The University of Queensland, Brisbane 4072, Australia; Australian Regenerative Medicine Institute, Monash University Clayton campus, Melbourne 3800, Australia; Department of Neurology, University Hospital Hamburg-Eppendor, Martinistr, 52, 20246 Hamburg, Germany; Queensland Institute for Medical Research, Herston, Brisbane 4006, Australia; Department of Pathology and Department of Neurology, National Taiwan University Hospital and National Taiwan University College of Medicine, Taipei, Taiwan; Institute of Neuroscience & Psychology, Wellcome Surgical Institute, Garscube Estate, University of Glasgow, Glasgow G61 1QH, United Kingdom; School of Dentistry, The University of Queensland, Brisbane 4072, Australia; Stroke Center, Department of Neurology, National Taiwan University Hospital, Taipei, Taiwan; The Jackson Laboratory, Bar Harbor, ME 04609, United States of America; Queensland Brain Institute, The University of Queensland, Brisbane 4072, Australia.; Faculty of Medicine, Department of Anatomy and Neuroscience, The University of Melbourne, 3010, Australia.

**Keywords:** C-type lectin, ischemia, MCAO, microglia, sterile inflammation.

## Abstract

Mincle is a C-type lectin known to play a role in innate immune responses to sterile inflammation, but its contribution to pathologies following an ischemic or traumatic injury is not well understood. In the current study we demonstrate a key role for Mincle in ischemic (i.e. transient middle cerebral artery occlusion) but not traumatic central nervous system injury; absence of Mincle also did not significantly alter the extent of tissue damage or functional outcome in peripheral models of ischemic tissue injury. In the stroke model mice lacking Mincle displayed significantly improved functional outcome from focal cerebral ischemia. The functional improvements in Mincle KO animals were accompanied by reduced infiltration of neutrophils and lower levels of proinflammatory cytokines in recruited peripheral blood cells. Bone marrow chimera experiments revealed that presence of Mincle in the central nervous system, but not peripheral immune cells, was the critical regulator of a poor outcome following transient focal cerebral ischemia, however we exclude a direct role for Mincle in microglia or neural activation. We demonstrate that Mincle lacks widespread expression in the brain, but is specifically associated with macrophages resident in the perivascular niche. These findings implicate Mincle in the initiation, extent and severity of local responses to ischemic injury in the brain, but not peripheral tissues. Mincle signalling therefore offers a novel therapeutic target in the quest to limit damage after stroke.

**Sources of support:** Australian National Health & Medical Research Council [1057846, 1060538 and Fellowship to NAR], SpinalCure Australia (Career Development Fellowship to MJR), the Australian Research Council, the State Government of Victoria, the Australian Government and The University of Queensland.

## Introduction

Ischemic stroke results in the damage and death of neurons in the perfusion territory of the affected blood vessel; the neurodegenerative mechanisms involve metabolic and oxidative stress, excitotoxicity and apoptosis, as well as neuroinflammation and the infiltration of activated leukocytes^1^. Chronic inflammatory conditions including arteriosclerosis, obesity and infection, also increase the risk of stroke and worsen outcome^2,3^. Sterile inflammation is therefore an important clinical target in tissue injury arising from cerebral ischemia. Molecular models of innate immune signalling suggest that excessive / deregulated inflammation is a confounding factor, in stroke as well as other forms of central nervous system (CNS) trauma. For example, mouse knock-out models focusing on innate immune receptors, such as the Toll-like receptor (TLR) family, suggest that blocking inflammation is beneficial^4,5^ to limit tissue damage. In contrast, targeting the TLR adaptor protein, MyD88 provided no benefit for neurological outcomes and worsened neuronal cell death in animals subjected to global or focal ischemic injuries^6,7^. Likewise, in mouse models of spinal cord injury (SCI), blocking Tlr2 or Tlr4 reduces microglia and/or astrocyte activation^8^ but worsens tissue damage and functional recovery^9^. The presence of TLRs in the central nervous system indicates pleiotropic, possibly neuroprotective roles in sterile injury, whereas increased expression of TLR2 and 4 in peripheral leukocytes is concordant with higher inflammatory markers in clinical stroke, and is predictive of poor outcome in some stroke patients^10^.

Clinical studies have shown that high circulating neutrophil numbers or high neutrophil to lymphocyte ratio are positive predictors of stroke severity^11^. Strategies that reduce neutrophil influx to the site of ischemic injury reduce inflammation, collateral blood vessel occlusion and the severity of injury^12–14^. In contrast, blocking the phagocytic clearance of dead cells by microglia will exacerbate injury^15^, and reparative roles of tissue macrophages are necessary for wound healing and functional recovery from sterile inflammation^16^. It is thus becoming increasingly apparent that functional differences in infiltrating myeloid cells (e.g. inflammatory neutrophils vs. monocytes) during the acute phase of sterile injury are important determinants of neuroprotection, blood brain barrier integrity, and recovery from stroke.

Innate immune cells, particularly macrophages and neutrophils, express a variety of receptors for endogenous ligands that are candidate regulators of inflammation during ischemic injury. The C-type lectin, Dectin-1, is a myeloid receptor that antagonises Tlr2 in SCI. Specifically, *Dectin-1* knock-out mice were protected from axonal dieback after traumatic injury, and in the wild type (WT) animals, Tlr2 activation reduced the harmful effects of Dectin-1 on axonal damage.^17^ Mincle (also designated as *Clec4e*) is a myeloid receptor closely related to Dectin-1 that has been reported to recognize necrotic cells via detection of the nuclear protein Sap130^18^. Mincle was originally identified as a lipopolysaccharide (LPS)-inducible protein in macrophages and has subsequently been shown to stimulate inflammatory responses to fungal and mycobacterial pathogens^19–22^. Mincle associates with the Fc receptor common gamma chain (FcR?) in immune cells, triggering intracellular signaling through the spleen tyrosine kinase (Syk) and the caspase recruitment domain protein Card9, which drives production of inflammatory cytokines such as TNF^23,24^; inhibition of Syk limits thrombosis and vascular inflammation in a variety of animal models of sterile injury^25^. Therefore, the Mincle/Syk axis is likely to also contribute to the pathophysiology of inflammation in ischemic stroke.

Two previous studies have suggested a deleterious role for Mincle in rodent models of subarachnoid haemorrhage^26^ and ischemic stroke^27^. Both used Syk inhibition, which is not selective for Mincle signalling, and as a result the role of Mincle in stroke outcomes remains poorly defined. In the current study, we used Mincle knockout mice (*Clec4e*^-/-^) to demonstrate that absence of Mincle, specifically in the central nervous system, significantly improves ischemic stroke outcomes. Mincle does not have the widespread brain expression previously described^26,27^, but instead appears restricted to perivascular macrophages and peripheral leukocytes. Absence of Mincle did not affect outcomes following traumatic spinal cord injury, or of ischemic injuries in other organs, such as the heart or the intestine. The combined data presented here suggest a key role for Mincle in ischemic CNS injuries where the integrity of the blood-brain/spinal cord barrier is not compromised by mechanical forces during the initiating event.

## Materials and Methods

### Animals and reagents

All experimental procedures followed the “Australian code of practice for the care and use of animals for scientific purposes”, and were approved by The University of Queensland and Monash University Animal Ethics Committees (ethics license numbers SBMS/358/12/NHMRC/ARC, SBMS/085/09, MARP-2011-175 and SBMS/311/12/SPINALCURE). The mouse colony was maintained in conventional or specific pathogen free conditions but, leading up to experiments, all animals spent at least one week in conventional housing conditions, with autoclaved sawdust or corn cob for bedding and a maximum of 5 mice per cage. Mice were kept in a conventional light cycle (12h light/12h dark), with controlled temperature (22-26 °C) and humidity (40-60%), and had access to normal chow diet and water *ad libitum*. Autoclaved cardboard boxes were used as environmental enrichment, and animals were checked for health daily. Homozygous null C57Bl/6J *Clec4e*^-/-^ mice were used as previously described^22^, and they were compared to either WT C57BL/6J or cohoused *Clec4e*^+/-^ littermates, which have been shown to display the same immune phenotype as WT C57BL/6J mice^28^.

Four different laboratories conducted the surgeries described herein. In all cases, except where indicated, a randomized experimental design consisted of pre-assigned groups of mice where the surgeon or operator was blind to genotype.

All genotyping was conducted by PCR, using primers: *ON14*, ATTGCCACTGACCCTCCACC; *MN469*: CCCCTGTCACTGTTTCTCTGCA; *MN473*: TGCAGCCCAAGCTGATCCTC. The Syk inhibitor BAY61-3606 (B9685, Sigma-Aldrich) was injected in the femoral vein at a dose of 1mg/kg (25 μl), 30 min before middle cerebral artery occlusion (MCAO), or 3 h post-reperfusion. All experiments took place in physical containment 2 (PC2) laboratories. All sections of this study adhere to the ARRIVE Guidelines for reporting animal research (Supplementary Figure 2).

### Focal cerebral ischemia model

Male, 3 to 6 month-old mice were anesthetized for focal cerebral ischemia by transient middle cerebral artery occlusion (tMCAO). Male mice were individually housed. The first round of surgery (Figure 1 and Figure 2) did not use a randomized experimental design. For all other tMCAO, including bone marrow chimeras (Figure 3, n=70), and microglia profiling (Figure 5, n=16), operators were blinded to genotype or treatment group and randomization was based on predesigned lists using colour coded cages and reagents. Exclusion criteria were excessive bleeding or death within 24 h after tMCAO. On these grounds, 1 out of 10 Syk inhibitor pre-treated, 2 out of 16 *Clec4e*^-/-^ and 7 out of 34 WT animals with tMCAO were excluded. Mice were anesthetized with 2% isoflurane in oxygen with spontaneous breathing and body temperature at 37°C. After a midline neck incision, the left external carotid and pterygopalatine arteries were isolated and ligated with 5-0 silk thread. The internal carotid artery was occluded with a clip at the peripheral site of the bifurcation to the pterygopalatine artery and the common carotid artery was then ligated with 5-0 silk thread. The external carotid artery was cut and a 6-0 nylon suture with a blunted tip (0.20 mm) was inserted. The clip at the internal carotid artery was then removed for advancement of the nylon suture into the middle cerebral artery to slightly more than 6 mm from the internal carotid-pterygopalatine artery bifurcation. After 1 h occlusion, the nylon suture and ligatures were removed to initiate reperfusion for 24 h up to 7 days. In the sham group, these arteries were visualized but not disturbed. Animals were subjected to cerebral blood flow (CBF) measurements using a laser Doppler perfusion monitor (Moor Lab) to confirm MCAO. The Doppler laser tip was placed perpendicular to the surface of the right parietal skull (1 mm posterior and 5 mm lateral to the bregma) to monitor blood flow in the middle cerebral artery territory.

**Figure 1.**
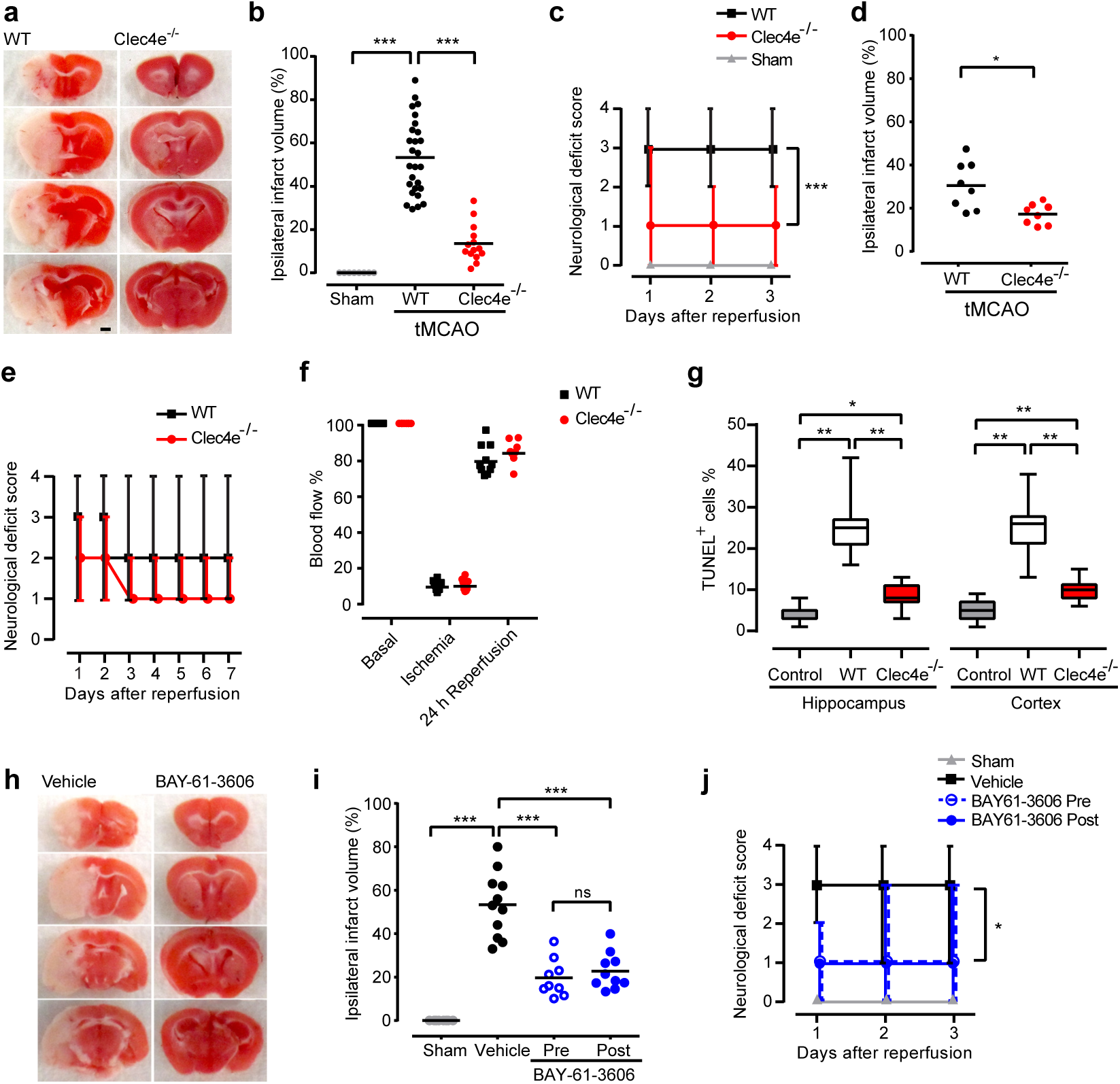
Mincle signaling worsens brain damage and functional outcome after transient MCAO. Mice of the indicated genotypes were subjected to sham surgery or transient middle cerebral artery occlusion (tMCAO), and brain damage and neurological function were evaluated (a) Representative TTC-stained brain sections from WT and *Clec4e*^-/-^ mice 3 days post-reperfusion. Infarct volumes (b) for WT mice subjected to sham surgery (n = 6), and WT (n=27) and *Clec4e*^-/-^ (n=14) mice 3 days after tMCAO. Bars represent mean. (c) The corresponding daily neurological deficit scores are shown as median and range. (d) Infarct volumes for WT (n=8) and *Clec4e*^-/-^ (n=8) mice 7 days after tMCAO are significantly different as shown by t-test. (e) The corresponding daily neurological deficit scores are shown as grouped with median and range. (f) Laser Doppler flowmetry shows no differences between samples in the extent to which MCAO compromises blood flow. (g) TUNEL positive cells were quantified in both hippocampus and cortex from control (n=4), WT (n=6) and *Clec4e*^-/-^ (n=6) mice after global cerebral ischemia. The percentage of TUNEL positive cells is represented showing median, the 25th to 75th percentiles, and min-max range. (h) Representative TTC-stained brain sections from mice treated with vehicle or the Syk inhibitor BAY-61-3606, 3 days post-reperfusion. (i) Infarct volumes for WT mice subjected to sham surgery (n = 6), vehicle-treated (n = 11) mice, or mice treated with BAY-61-3606 before MCAO (n = 9) or 3 h after the onset of reperfusion (n = 10), 3 days after tMCAO. (j) The corresponding daily neurological deficit scores are as median and range, and both pre-and post-treated samples are significantly different from vehicle control. Tests: (b,I,g) ANOVA, (c,j) Kruskal-Wallis, (d) t-test, (e) Mann-Whitney. ***: p < 0.001, **: P<0.01, *: P<0.05, ns: not significant; comparisons indicated by brackets. Scale bar for images: 1 mm.

**Figure 2.**
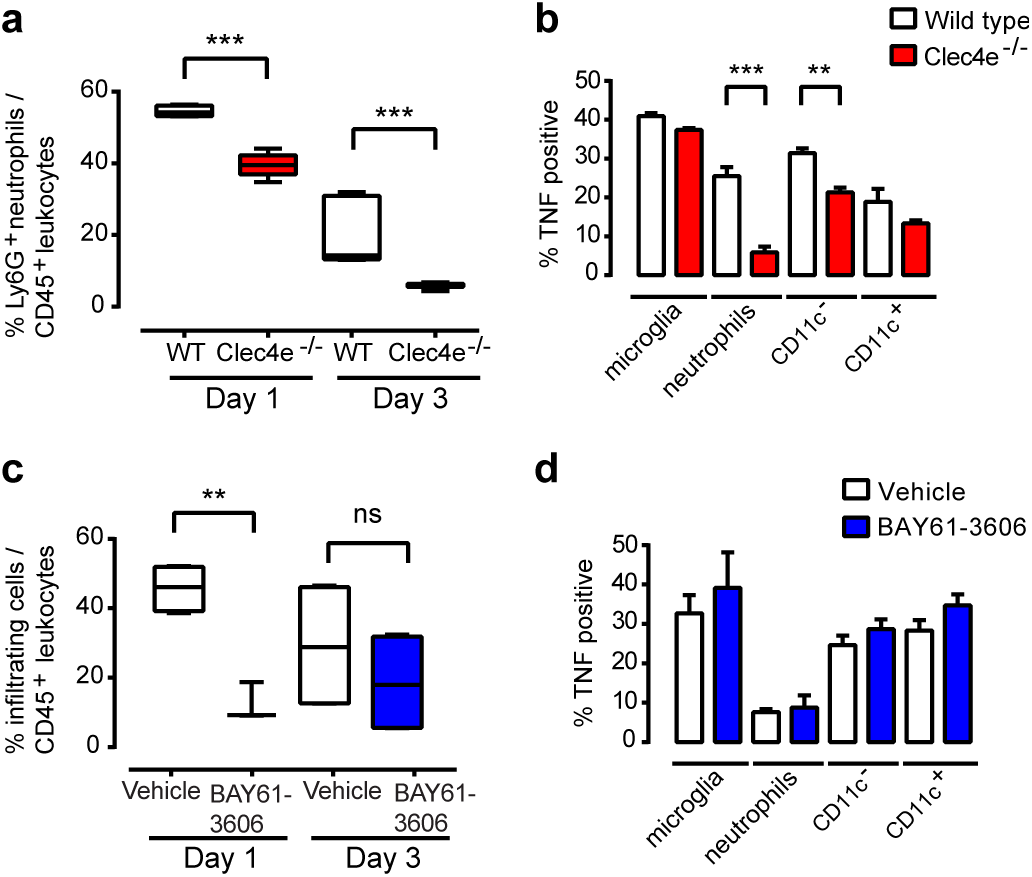
Neutrophil infiltration was significantly reduced in *Clec4e*^-/-^ mice following tMCAO. (a) Flow cytometry of leukocytes in the ipsilateral hemisphere showed a significantly lower proportion of infiltrating neutrophils (CD45^high^, CD11b^+^, Ly6G^+^) in *Clec4e*^-/-^ mice at 1 and 3 days post-reperfusion (box and whiskers plot). (b) There were fewer TNF-positive infiltrating neutrophils and CD11c^-^ monocytes in *Clec4e*^-/-^compared to WT mice 24 h after reperfusion. (c) Flow cytometry revealed a significantly lower proportion of infiltrating leukocytes (CD45^high^, CD11b^+^) in *Clec4e*^-/-^mice than in WT controls 24 h post-reperfusion. (d) No differences in the proportion of infiltrating neutrophils or in TNF-positive leukocytes were observed. t-test: ***: P < 0.001, **: P < 0.01, ns: not significant. Comparisons indicated by brackets. % cells in (a) and (c) are represented by box and whiskers plots, showing median, the 25th to 75th percentiles, and min-max range. % TNF-positive cells are represented by mean + s.e.m.

### Quantification of cerebral infarction and neurological deficit assessment

At 3 or 7 days post-reperfusion, mice were euthanized by an investigator blinded to the genotype and/or treatment, the brains removed into PBS (4°C) for 15 min, and 2 mm coronal sections were obtained. These were stained with 2% 2,3,5-triphenyltetrazolium chloride (TTC, T8877 Sigma-Aldrich) at 37°C for 15 min. The stained sections were photographed, and the images digitized. The infarct area of each section was defined as the pale area surrounded by red undamaged tissue. Measurements were done using NIH image 6.1 software (National Institutes of Health, Bethesda, MD, USA). To correct for brain swelling, the infarct area was determined by subtracting the area of undamaged tissue in the left (ipsilateral) hemisphere from that of the whole contralateral hemisphere. Infarct volume was calculated by integration of infarct areas for all slices of each brain, and then expressed as a % of the ipsilateral hemisphere. The functional consequences of tMCAO were evaluated using a 5-point neurological deficit score (0, no deficit; 1, failure to extend right paw; 2, circling to the right; 3, falling to the right; and 4, unable to walk spontaneously^29^).

### Global ischemia model and TUNEL assay

Adult male, age-matched C57BL6/J WT (n=4) and *Clec4e*^-/-^ mice (n=6) were used for this investigation, and the investigators were not blinded to genotype. Transient global cerebral ischemia was performed by the 2-vessel occlusion model^1^. The common carotid arteries were occluded for 30 minutes and the blood was allowed to reperfuse the tissue for 72 h before euthanasia. Mice were euthanized, brains were fixed and embedded in paraffin wax, and sections were pretreated with 20 μg/ml proteinase K (Roche, Switzerland) in 10 mM Tris, pH 7.4. The In Situ Cell Death Detection Kit, POD (11684817910, Roche, Switzerland) was used according to manufacturer’s instructions, and tissue was counterstained with Gills haematoxylin. The area of positive (brown) cells was divided by the total section area to calculate the % TUNEL^+^ cells. Sections treated with 100 U/ml recombinant DNase I were used as positive control.

### Flow cytometry

Animals were euthanized and perfused with PBS prior to isolation of as described in ^30^. Brains were dissected, cerebella removed, and the left ischemic (ipsilesional) hemispheres selected. Three hemispheres were pooled, digested for 30 min at 37°C (1 mg/ml collagenase, 0.1 mg/ml DNAse I in DMEM), and pressed through a cell strainer (40 μm; BD Biosciences). Cells were incubated with standard erythrocyte lysis buffer on ice, separated from myelin and debris by Percoll (17-5445-01, GE Healthcare) gradient centrifugation, and then incubated with the following antibody cocktail for 30 min at room temperature in buffer (0.5% bovine serum albumin, 0.02% sodium azide in PBS): CD45 (25-0451-82, eBioscience, 1:100), Ly6G (560603, BD Biosciences, 1:100), CD11b (557396, BD, 1:300), CD11c (17-0114-82, eBioscience, 1:100) and TNF (554419, BD, 1:100). Data were acquired with a LSR II FACS system (BD Biosciences) and analyzed with FlowJo (TreeStar). Doublets were excluded with FSC-A and FSC-H linearity.

### Generation of bone marrow chimeric mice

Recipient mice (3-6 months of age) were sub-lethally irradiated with 2 doses of 5 Gy delivered 14 h apart and their immune system rescued via bone marrow (BM) transplantation from either WT or *Clec4e*^-/-^donors, 3-4 h after the second dose of irradiation. BM chimeric mice were left to recover for at least 3 months to allow full reconstitution of the peripheral immune compartment. Flow cytometric analysis confirmed over 95% donor engraftment efficiency based on the relative frequency of donor and host white blood cells, which were identified on the basis of CD45.1^+^/ CD45.2^+^ antigen expression. Mice were colour coded prior to surgery and then subjected to tMCAO by an experimenter blinded to the genotype or treatment groups, and infarct size was determined by TTC staining as detailed above.

### Spinal cord injury and assessment of locomotor recovery

Adult, age-and weight-matched female C57BL6/J WT (n=12) and *Clec4e*^-/-^ mice (n=14) were used for these experiments. Order of surgery was randomized based on predesigned lists, with the experimenter conducting the surgery remaining blinded to genotype throughout all aspects of surgery. In brief, mice were anesthetized via intraperitoneal injection with Xylazine (10 mg/kg, Ilium) and Zolazepam (50 mg/kg, Virbac) and subjected to a severe contusive SCI. The ninth thoracic (T9) vertebra was identified as described previously^31^, followed by a dorsal laminectomy as described previously^32,33^. This exposed the dorsal surface of the spinal cord at spinal level T11, where a force-controlled 70 kilodyne (kd) impact was applied using the Infinite Horizon impactor device (Precision Systems and Instrumentation). Paravertebral muscles were sutured post-impact using 5-0 Coated Vicryl (Polyglactin 910) sutures (Ethicon), followed by wound closure using Michel wound clips (KLS Martin Group). SCI mice were then randomly re-assigned to cages labelled A, B, C, etc., with 2-3 mice being housed per box. Post-operative care involved sub-cutaneous administration of a single dose of buprenorphine (0.05 mg/kg) in Hartmann’s Sodium Lactate solution for analgesia.

Additionally, a prophylactic dose of Gentamicin, (1.0 mg/kg, Ilium) was administered daily for 5 days post-injury. Recovery of hind-limb function was assessed using the 10-point Basso Mouse Scale (BMS), a system designed specifically for the assessment of murine locomotor recovery following SCI^34^. Multiple aspects of locomotion, including ankle movement, stepping, co-ordination, paw placement, trunk stability and tail position, were assessed using this scale. Experimental animals were randomly picked up from their cages and assessed by two investigators blinded to the genotype at 1, 4, 7 days post-SCI and then weekly thereafter up until the study endpoint (35 days post-injury). Animals with a deviation in more than ±5 kdyne from the mean force, or a spinal cord tissue displacement ±100 μm from the experimental mean were excluded from the study. Based on these criteria, 2 WT and 4 *Clec4e*^-/-^ mice were excluded from the study. For the remaining n=10 mice per genotype, the actual applied force and displacement for WT and *Clec4e*^-/-^ animals was 75.40±0.91 *vs*. 74.10±0.99 kdyne (p>0.34), and 537.7±15.22 *vs*. 535.9±18.11 μm, respectively (p>0.94). There was thus no bias in the severity of injury between genotypes at the outset.

### Spinal cord tissue sectioning and immunofluorescence

All mice were euthanized at 35 days post-injury. In brief, mice were deeply anesthetized using sodium pentobarbital and transcardially perfused with 15ml of saline (0.9% NaCl containing 2IU/ml Heparin (Pfizer) and 2% NaNO_2_), followed by 30ml of phosphate-buffered Zamboni’s fixative (2% Picric acid, 2% Formaldehyde, pH 7.2-7.4). Vertebral columns were excised and post-fixed overnight at 4°C. The spinal cord was then dissected and placed in sequential overnight incubations of 10% and 30% sucrose in PBS for cryoprotection, followed by embedding in Tissue-Tek Optimal Cooling Temperature (Sakura Finetek), and snap-freezing on dry-ice cooled isopentane. Transverse 20 μm thick sections of spinal cord were cut using a Leica Cryostat CM3050-S and collected in 1:5 series on Superfrost Plus slides. Sections were incubated with IHC blocking buffer (2% Bovine Serum Albumin and 0.2% Triton X-100) for one hour at room temperature (RT) in a humidified chamber, and then overnight at 4°C with primary antibodies: 1:1600 chicken anti-mouse GFAP (Abcam; #ab4674) and 1:200 rabbit anti-mouse fibronectin (Sigma-Aldrich; #F3648), or 1:1000 rabbit anti-mouse GFAP (Dako; #Z0334) in the absence of fibronectin staining. After washing, slides were incubated for 1 h at RT with the following secondary antibodies as required: 1:400 goat anti-chicken 555 (Abcam; #ab150170) and 1:400 goat anti-rabbit 488 (Thermo Fisher Scientific; #A-11034) and 1:150 FluoroMyelin Red (Thermo Fisher Scientific; #F34652). Hoechst 33342 nuclear dye was used for counterstaining. After washing and mounting the slides, images were captured on a single plane using a Zeiss Axio Imager and Zen Blue 2012 Software (Zeiss). ImageJ software (National Institutes of Health) was utilized for analysis. Section areas were determined by outlining the section boundary on the GFAP^+^ channel (excluding the leptomeninges). Proportional area measurements were calculated by thresholding the FluoroMyelin Red stained area in ImageJ and dividing it by the total section area. Lesion volumes and/or length were calculated by multiplying fibronectin^+^ areas by the section thickness and 1:5 series count.

### Intestinal ischemia and reperfusion, histological analysis and myeloperoxidase (MPO) quantification

Mice (WT, n = 11; *Clec4e*^-/-^, n = 10) were anesthetized with 2% isoflurane in oxygen through a facemask, with spontaneous breathing and body temperature at 37°C. The surgeries were not randomized, but tissues were collected into coded tubes and the analysis performed by an operator blind to genotype. A midline incision was made through the skin and then along the linea alba separating the rectus abdominis muscle. The exposed intestines were displaced and a ligature was tied with silk suture material around the superior mesenteric artery except in animals undergoing sham surgery. After 30 min of ischemia the ligature was removed, and after 2 h of reperfusion the mice were euthanized. For histological analysis, three portions of small intestine were stored in 4% paraformaldehyde for 24 h. Tissues were embedded in paraffin wax, sectioned transversely and stained with haematoxylin/eosin. The average of villi damage was determined after grading each of 100 villi per mouse on a 0–6 scale as previously described^35^. For MPO activity, three portions of small intestine were homogenized in 50 mM potassium phosphate, centrifuged and pellets resuspended in 0.25 mM hexadecyltrimethylammonium bromide (H5882, Sigma-Aldrich) for MPO solubilisation. After homogenisation and centrifugation, supernatants were assayed with 1.21 mg/ml o-dianisidine dihydrochloride (D3252, Sigma-Aldrich) and 2.17% hydrogen peroxide, and absorbance read at 460 nm.

### Myocardial infarction and echocardiography analysis

All animals in the same cage (siblings) were experimented on in a blinded fashion (for both echocardiography and surgery). Genotypes were checked after the experiment was finalized. To induce myocardial infarction, the left coronary artery (LDCA) of 10 week-old mice was ligated. For this procedure, animals were anaesthetized using 2% isoflurane and subjected to artificial ventilation through endotracheal cannulation. An incision was made through the muscle of the 4^th^ and 5^th^ intercostal space, and an 8-0 polyethylene suture passed under and tied around the LDCA 1 mm below the tip of the left auricle. Buprenorphine analgesic solution was administered subcutaneously (0.05 μg/g) twice a day for 3 days following surgery. For non-invasive echocardiography, control (heterozygous or WT) and mutant (homozygous) adult mice, in homeostasis or 1 month after surgical intervention, were anaesthetized (isoflurane) and kept sedated under 1.5% isoflurane. Imaging was performed in spontaneously breathing animals in prone position using Vevo 2100 Imaging System (FUJIFILM VisualSonics) equipped with 18 to 38 MHz linear array transducer. Standard parasternal long-and short-axis views were obtained to assess left ventricular chamber function. Breathing was also monitored to avoid measurement distortions during breathing cycle. Calculations of cardiac function were done using Vevo2100 Cardiac Measurements Package. Mice were euthanized one month after surgery, their hearts dissected and prepared for histology.

### Cell culture, and oxygen and glucose deprivation

Neuronal cultures were established from littermate 16 day-old WT, *Clec4e*^+/-^ or *Clec4e*^-/-^ mouse embryos. Genotypes were checked after the experiment was finalized. Cells were maintained at 37 °C in Neurobasal medium containing Glutamax and B-27 supplements, and 0.001% gentamycin sulfate (all Life Technologies). Cells were ascertained by immunofluorescence to be 95% neurons and 5% astrocytes, with the occasional microglia. Glial cultures were established from postnatal day 1 WT, Clec4e^+/-^ or Clec4e^-/-^ mice and seeded in DMEM/F12 medium containing Glutamax and gentamycin 10 mg/L (all Life Technologies), and 10% fetal bovine serum (FBS). Microglia were separated from astrocytes with the use of CD11b (Microglia) MicroBeads (Miltenyi Biotec). The murine brain endothelial cell line bEnd.3 (ATCC CRL-2299) was grown to confluence in DMEM supplemented with Glutamax (Life Technologies). For oxygen and glucose deprivation (OGD), cultures were incubated with glucose-free Locke’s buffer (in mmol/L: 154 NaCl, 5.6 KCl, 2.3 CaCl_2_, 1 MgCl_2_, 3.6 NaHCO_3_, 5 HEPES, pH 7.2, supplemented with gentamycin 10 mg/L), placed in an incubator where the oxygen was displaced with nitrogen to a level of 0.2%, and incubated for 3 hours. Incubation with trehalose dimycolate (Sigma Aldrich) was conducted for 24 hours.

### Immunofluorescence

For analysis of Mincle expression, primary microglia and RAW264.7 cells (ATCC TIB-71) were grown on 12 mm microscope coverslips and fixed in 4% paraformaldehyde. Primary antibodies used for staining were rat anti-Mincle (clone 1B6, D266-3M2, MBL) and goat anti-ionized calcium-binding adapter molecule 1 (Iba1, polyclonal, ab5076, Abcam). For immunofluorescent staining on rat tissue, 6 μm microtome sections from Wistar Kyoto or spontaneously hypertensive stroke-prone (SHRSP) rats subjected to permanent MCAO for 24 h using diathermy with modification were obtained from a previous study^36^. Primary antibodies used were rat anti-Mincle (clone 4A9, D292-3M2, MBL), mouse anti-Mincle (clone 16E3, ab100846, Abcam); mouse anti-alpha smooth muscle actin antibody (alpha-SMA, clone 1A4, ab7817, Abcam); rabbit anti-glial fibrillary acidic protein (GFAP, polyclonal, ab4674, Abcam), and mouse anti-CD163 (clone ED2, Santa Cruz Biotechnology, sc-59865). Secondary antibodies were conjugated with Alexa Fluor 488, 568 and 647 (Life Technologies). Hoechst 33342 nuclear dye was used for counterstaining. Images were acquired using an Olympus BX61 microscope (Japan).

### qRT-PCR

For qPCR, RNA was isolated with RNeasy Plus Mini Kit (74134, Qiagen), its quality assessed using Nanodrop, and cDNA synthesized using iScript Reverse Transcription Supermix for RT-qPCR (Bio-Rad). qPCR was performed with FastStart Universal SYBR Green Master [Rox] (04913850001, Roche) in a C1000 Thermal Cycler Chassis with CFX96 Optical Reaction Module (Bio-Rad). The following Clec4e primers were used: *forward*, TGCTACAGTGAGGCATCAGG; *reverse*, GGTTTTGTGCGAAAAAGGAA. MIP2a primers were: *forward*, GAGACGGGTATCCCTTCGAC; *reverse*, TTCAGGGTCAAGGCAAACTT.

### Microglia isolation and microarray

WT and *Clec4e*^-/-^ mice were coded and randomized for both surgery and FACs profiling as described above. After 1 h tMCAO (as described previously) and 24 h reperfusion, mice were perfused with PBS, their brains dissected, and 2 ipsilesional hemispheres (with cerebellum and brainstem removed) pooled for microglia isolation. For sham-operated animals, the whole forebrain was used and brains were not pooled. Tissue was minced with a razor blade, triturated by pipetting up and down gently 20 times, and pressed through a 40 μm cell strainer; all steps were carried out on ice. After myelin separation by Percoll gradient centrifugation, around 80,000 CD45^intermediate^, CD11b^+^ microglial cells were sorted from each sample. Doublets were excluded with FSC-A and FSC-H linearity, and dead cells excluded using Zombie Violet^™^ Fixable Viability Kit (423113, BioLegend). RNA was isolated with RNeasy Micro Kit (74004, Qiagen), and yield and quality measured with the RNA 6000 Pico Kit (Agilent Technologies, 5067-1513) for Agilent Bioanaliser. Yield ranged between 1.4 and 11 ng, and the RNA integrity number ranged between 8.1 and 10. Samples were amplified with the GeneChip WT Pico Kit (902623, Affymetrix) and processed with the Mouse 2.0ST Gene Array WT pico assay (902463, Affymetrix). The last three steps were carried out by the Ramaciotti Centre for Genomics, University of New South Wales. The expression data (RMA background corrected, quantile normalized) is hosted in the www.stemformatics.org resource (dataset S4M-6731)^37^ and is available for download from GEO (Accession GSE77986). The expression threshold (detection floor) was calculated as the median expression of all antigenomic probesets on the microarray resulting in a value of log_2_ 3.38. The median of expression was taken to be the median of above-threshold log_2_ normalized probe expression values, which was log_2_ 6. Probes that failed to be expressed above detection threshold log_2_ 3.38 in the majority of biological replicates in at least one comparison group were removed from the analysis. The R/Bioconductor software package *limma*^38^ was used to find differentially expressed genes (DEG), with a false discovery rate threshold value of 0.01. Volcano plots were generated to visually inspect DEG significance vs. fold change.

### Data analysis

Unless indicated otherwise, the experimental unit for *in vivo* experiments was a single animal. The overall significance of the data was examined by one-way analysis of variance (ANOVA) followed by Neuman-Keuls multiple comparisons test, or two-tailed Student t-test, as appropriate. The differences between the groups were considered significant at P<0.05. Neurological deficit scores were analyzed by using a nonparametric Kruskal–Wallis test followed by Dunn’s multiple comparison test, or Mann-Whitney test. Physiological function and disease related studies require a good number of specimens to be considered statistically significant, due to intrinsic variations between individuals. In all mouse models tested, morphological and physiological differences may result in 15-25% variation in the severity of the surgically induced injuries. The initial surgical cohort sizes were determined by effect sizes observed with historical datasets, and observed variation within experimental groups was consistent with historical datasets. Unless stated, error bars show standard error of the mean (s.e.m.).

## Results

### Mincle deficiency improves functional outcomes and reduces infarct size in mouse models of cerebral ischemia

To directly address a role for Mincle in stroke-induced inflammation and tissue damage, we examined *Clec4e*^-/-^ mice and isogenic controls after 1 h of transient middle cerebral artery occlusion (tMCAO) and reperfusion. On average, *Clec4e*^-/-^ mice showed a 50% reduction in infarct volume as assessed by TTC staining at 3 days post-stroke (Figure 1a,b). *Clec4e*^-/-^ mice also had significantly lower neurological deficit scores than WT controls (Figure 1c). These findings were confirmed and corroborated upon in an independent experiment in which mice were monitored up to 7 days post-stroke, with differences in infarct size and neurological deficit scores continuing to be significantly different between groups (Figure 1d,e). Improved outcomes in *Clec4e*^-/-^ mice could not be explained by the effect of MCAO on arterial blood flow, as laser Doppler flowmetry revealed similar reductions in blood flow during ischemia, and a comparable recovery 24 h after reperfusion (Figure 1f). The benefits of *Clec4e* deletion were also confirmed in another model of ischemia, i.e. global cerebral ischemia. Here, *Clec4e*^-/-^ mice showed significantly fewer TUNEL-positive cells in hippocampus and cortex (Figure 1g), indicating an effect of Mincle in the extent of apoptosis induced by ischemia.

As the kinase Syk mediates Mincle signaling in macrophage responses to pathogens^20^, we next used the Syk inhibitor BAY61-3606 to assess the effect of Syk in stroke-induced injury. Mice treated either before tMCAO or 3 h after reperfusion showed an over 50% decrease in infarct volume (Figure 1h,i) and a better functional outcome compared to vehicle-treated mice (Figure 1j), supporting the potential efficacy of Syk inhibitors in this stroke model.

### Mincle and its downstream partner Syk mediate reperfusion-induced leukocyte brain infiltration

To determine the role of Mincle in inflammatory cell infiltration after tMCAO, immune cell populations present in ipsilateral brain hemispheres from WT and *Clec4e*^-/-^ mice were analysed by flow cytometry over the first 3 days post-reperfusion. The recruitment of neutrophils (CD45^high^, CD11b^+^, Ly6G^+^) to the injured brain parenchyma at 1 or 3 days after tMCAO was significantly attenuated in *Clec4e*^-/-^ mice (Figure 2a). In addition, the percentage of TNF-positive cells 1 day after reperfusion was significantly reduced in *Clec4e*^-/-^ neutrophils and CD11c^-^ monocytes (CD45^high^, CD11b^+^, CD11c^-^) (Figure 2b). It is noteworthy that *Clec4e*^-/-^ microglia (CD45^intermediate^, CD11b^+^), which appeared to be the primary source of TNF in the injured hemisphere, showed no difference in TNF presence between genotypes (Figure 2b). A single dose of the Syk inhibitor 3 h after tMCAO also impacted on inflammatory cell recruitment, leading to a significant reduction in the total CD45^high^ infiltrate on day 1 (Figure 2c); vehicle and inhibitor-treated animals had equivalent numbers of infiltrating leukocytes on day 3 after tMCAO. Unlike *Clec4e*^-/-^ mice, animals treated with Syk inhibitor did not show differences in TNF-positive myeloid cells (Figure 2d).

### Ischemia-induced tissue damage is driven by Mincle expression in the brain, not peripheral blood cells

We next aimed to determine whether Mincle influences stroke outcomes via its expression in circulating myeloid cells, CNS resident cells, or a combination thereof. Bone marrow chimera experiments showed that a lack of Mincle in the peripheral immune compartment (*Clec4e^-/-^ > WT*) had a small but significant effect on infarct volume after tMCAO (Figure 3a,b), indicating a partial contribution of Mincle in bone marrow-derived cells to the *Clec4e*^-/-^ phenotype. In contrast, *Clec4e*^-/-^ mice that received a WT bone marrow transplant (*WT > Clec4e^-/-^*), which reinstates Mincle expression on circulating myeloid but not CNS-resident cells, were equally well protected against tMCAO as the chimeric group with global *Clec4e* deficiency (*Clec4e^-/-^ > Clec4e^-/^)* based on both infarct volumes and neurological deficit scores (Figure 3a,b). Loss of Mincle on CNS resident cells thus appears to be the main or critical contributor towards the protective phenotype observed in Clec4e^-/-^ mice.

**Figure 3.**
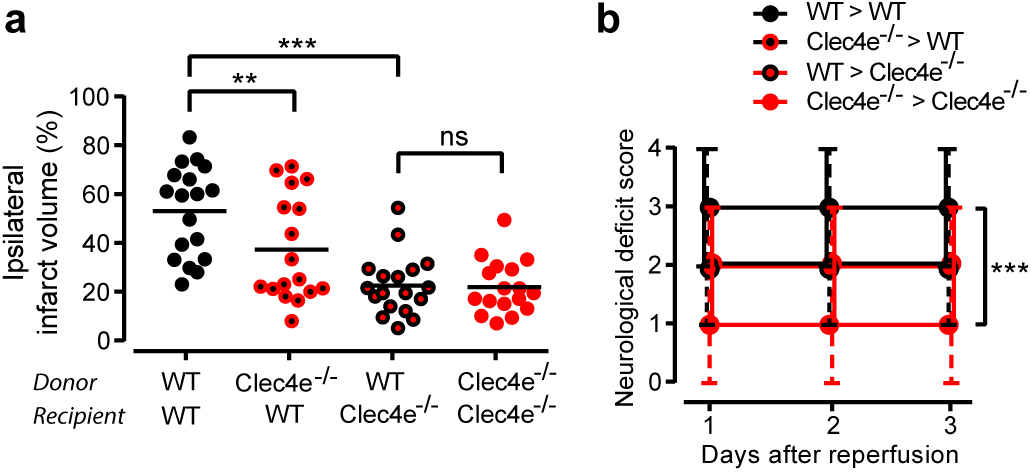
The origin of the *Clec4e*^-/-^ protective effect is in the central nervous system. (a) Ipsilateral infarct volume 3 days after tMCAO from bone marrow chimeras obtained between WT (C57BL/6J mice carrying the CD45.1 allele) and *Clec4e*^-/-^ animals. Donor and recipient genotypes indicated on the X-axis. Bars represent median and range. (b) Neurological deficit scores of bone marrow chimeras, assessed daily for 3 days, shown as median and range, where both Clec4e^-/-^ recipient samples are significantly different from WT>WT sample. WT donor to WT recipient, n=18; *Clec4e*^-/-^ donor to WT recipient, n=17; WT donor to *Clec4e*^-/-^ recipient, n=18; *Clec4e*^-/-^ donor to *Clec4e*^-/-^recipient, n=17. ANOVA: ***: P < 0.001, **: P < 0.01, ns: not significant. Comparisons indicated by brackets.

### Mincle deficiency does not influence the outcome from traumatic CNS injury or peripheral organ ischemia

We theorized that if Mincle is a general activator of inflammation in the local response to tissue damage, then it may also play a key role in other models of tissue injury with sterile inflammation. Using a model of blunt CNS trauma, we first assessed whether absence of Mincle influenced the outcome from contusive SCI. Lack of Mincle did not result in an altered and/or improved recovery from contusive SCI, with no differences observed between genotypes for hindlimb locomotor performance up to at least 35 days post-injury (P>0.05, Figure 4a,b). Consistent with the behavioural outcome, absence of Mincle did not affect lesion volume, lesion length, or myelin preservation (Figure 4c-g).

**Figure 4.**
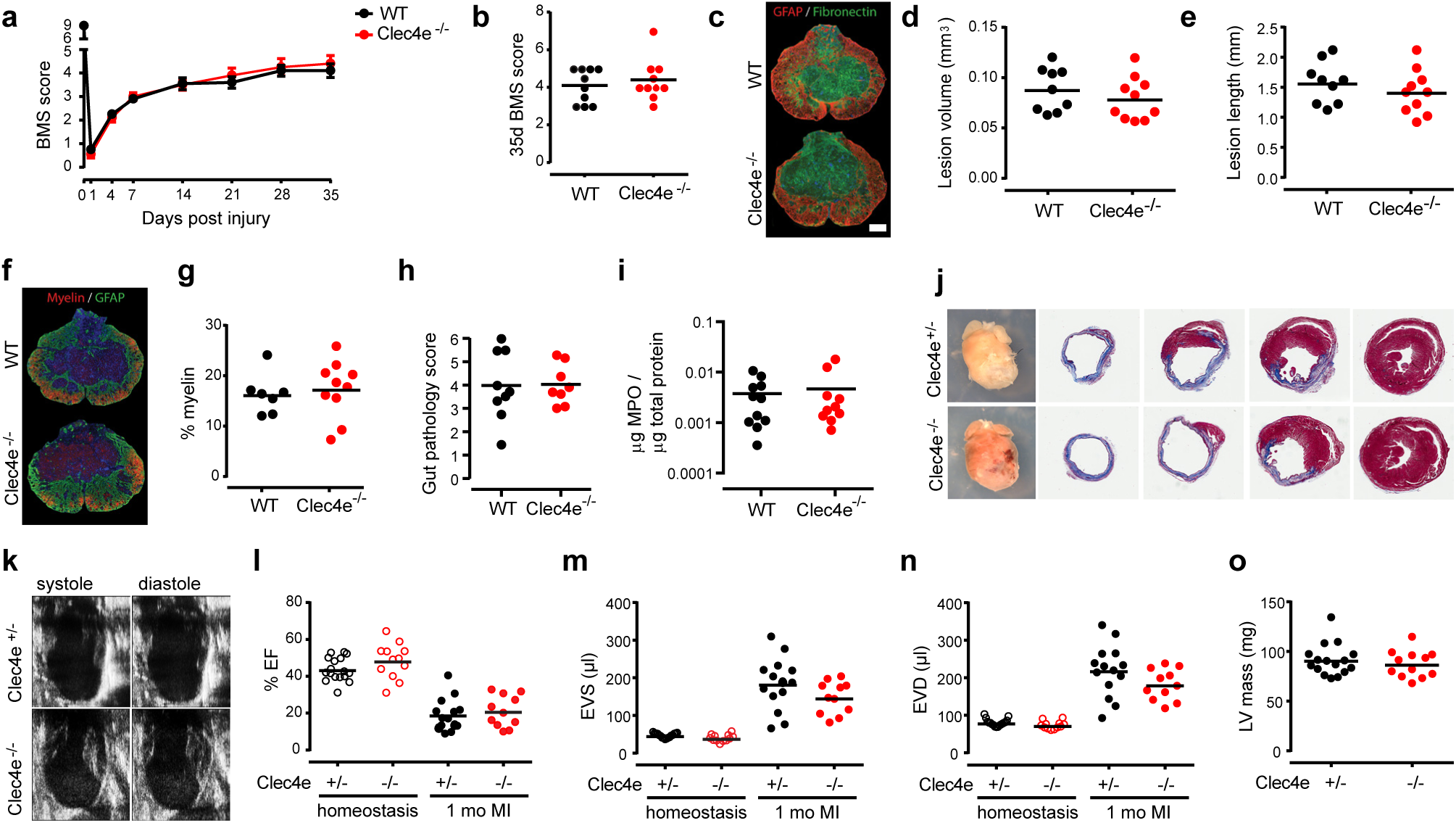
Absence of Mincle does not affect the outcome of spinal cord injury, intestinal ischemia and reperfusion (I/R) or myocardial ischemia. (a) No significant differences between genotypes (n=10) in 35 days post-injury BMS locomotor scores (t-test; P>0.05), or at endpoint (b). (c) Representative GFAP^+^/Fibronectin^+^ staining for the quantification of lesion core in WT and *Clec4e*^-/-^ mice. No significant differences were found between genotypes in either lesion core volume (d, t test P>0.05) or length (e, t-test P>0.05). Bars represent mean. Scale bar for images: 200 μm. Quantification of myelin staining in WT and *Clec4e*^-/-^ spinal cords (f) revealed no significant differences between genotypes (g). Intestinal I/R revealed no differences between genotypes with regards to pathology scores (h) or to levels of myeloperoxidase (MPO) an indicator of neutrophil infiltration (i). Whole mount views and histological sections of control and mutant hearts after 1 month of infarction (j). Trichrome staining, representative of n=10, shows scarring (blue) and viable myocardium (red) at various transverse levels of the heart, from apex (left panel) to base (right panel). (k-o) Echocardiography analysis of control and mutant hearts after 1 month of infarction (*Clec4e*^+/-^ & WT animals, n=14, *Clec4e*^-/-^ animals, n=11). (k) Representative images of infarcted hearts in systole and diastole. Ejection fraction (EF - l), end systolic volume (ESV – m) and end diastolic volume (EDV – n), as well as other analysed parameters, are not significantly altered in mutant hearts, suggesting that *Clec4e* is not implicated in the regenerative response after myocardial infarction. Left ventricular mass (LVM – o) was used as an indication of heart size in homeostasis and is not significantly different between control and mutant hearts.

Lack of Mincle also did not influence outcomes in two models of peripheral ischemic injury, namely intestinal ischemia and reperfusion, and myocardial infarction. Specifically, WT and *Clec4e*^-/-^ mice subjected to 30 min of ischemia and 2 h of blood reperfusion showed similar gut pathology scores (Figure 4h). Neutrophil recruitment to the injured intestine was also not changed in the absence of Mincle based on myeloperoxidase content in the tissue (Figure 4i). Following myocardial infarction, a similar level of scarring was observed between the hearts of *Clec4e*^-/-^ mice (n=11), *Clec4e*^+/-^ littermates and WT animals (combined n=14) at 1 month post-injury (Figure 4j). Echocardiography analysis of heart chamber function, either in homeostasis or 1 month after surgical intervention (Figure 4k), revealed no significant differences between genotypes in ejection fraction (Figure 4l), or systolic and diastolic end volume (Figure 4m,n). Left ventricular mass was also similar for both genotypes in homeostasis (Figure 4o), confirming a lack of size bias in baseline homeostatic functional parameters. Collectively, these data indicate that Mincle is not a determinant of outcomes in models of neurotrauma or peripheral ischemic tissue injury.

### Mincle expression in the brain is restricted to a specific cell type

To better understand the unique role of Mincle in ischemic stroke, we performed a careful search for which cell(s) in the CNS express Mincle. Given that Mincle is a myeloid cell receptor, we first assessed its expression in microglia. Immunofluorescence with 1B6, a rat antibody that recognises Mincle in the mouse macrophage cell line RAW264.7, showed no expression of Mincle in microglia cultures derived from neonatal brains. Stimulation with oxygen and glucose deprivation (OGD) as an *in vitro* model of ischemia also failed to reveal/upregulate Mincle expression (Figure 5a). We next determined whether microglia were sensitive to trehalose dimycolate (TDM) stimulation, a mycobacterial component known to signal through Mincle in macrophages, resulting in the production of the cytokine CXCL2 (MIP2a)^20^. Cultured microglia upregulated MIP2a expression in response to OGD regardless of their genotype, however WT (*Clec4e^+/-^*) microglia failed to induce MIP2a mRNA in response to TDM stimulation, indicating a lack of Mincle-dependant signalling in these cells (Figure 5b). *Clec4e*^+/-^ primary microglia did, however, upregulate *Clec4e* mRNA in response to OGD (Figure 5c), suggesting the possibility that Mincle could still affect the behaviour of adult microglia in tMCAO. We therefore sorted microglia (CD45^intermediate^, CD11b^+^) by FACS from the ipsilesional hemispheres of *Clec4e*^-/-^ and WT controls, 24 h after either tMCAO or sham surgery, for microarray profiling. *Clec4e* mRNA was not upregulated in WT microglia after tMCAO (Figure 5d). As expected, dramatic alterations in gene expression were observed between microglia from tMCAO mice compared to their sham-operated counterparts (Figure 5e), but the presence or absence of Mincle itself did not affect microglial gene expression (Figure 5f). This finding is in agreement with the observation of comparable TNF expression in WT and *Clec4e*^-/-^ microglia in earlier tMCAO experiments (Figure 2b). Mincle thus appears not to be expressed by microglia, nor does it play a direct role in acute microglia activation in *ex vivo* or *in vivo* ischemic stroke models. These observations thus exclude microglia as the primary mediators of the neuroprotective phenotype in stroked *Clec4e*^-/-^ mice.

**Figure 5.**
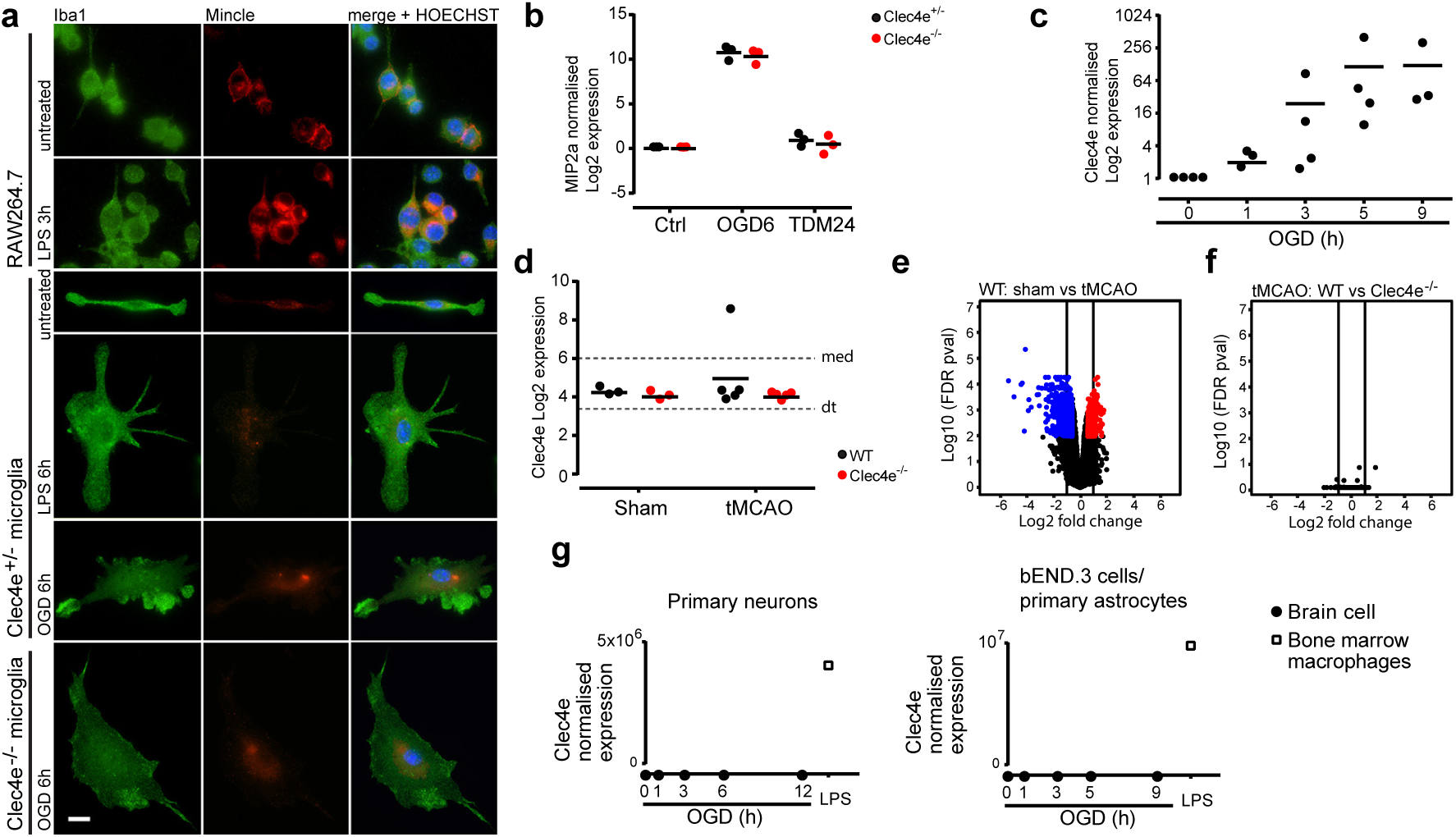
Mincle could not be found in microglia, neurons, astrocytes or endothelial cells. (a) An antibody that detects Mincle in the mouse macrophage cell line RAW264.7, and shows its upregulation by LPS, does not detect Mincle in cultured microglia in basal conditions or stimulated with LPS or oxygen and glucose deprivation (OGD). Scale bar: 10 μm. (b) MIP2a qRT-PCR in *Clec4e*^+/-^ and *Clec4e*^-/-^ cultured microglia from neonatal mice shows that Mincle does not contribute to *MIP2a* gene expression in microglia, and Mincle sufficient microglia do not produce *MIP2a* in response to the Mincle-specific ligand trehalose dimycolate (TDM). Data points represent different cultures, each derived from a single mouse (c) *Ex vivo* microglia upregulate *Clec4e* expression under OGD, as measured by qRT-PCR. (d) WT microglia sorted from the ipsilesional brain hemisphere 24 h after tMCAO do not show *Clec4e* upregulation, as shown by microarray, sham n=3 per genotype, tMCAO n=5 per genotype (dt: detection threshold, 3.38; med: median, 6) (e,f) Volcano plots for differences in microglial gene expression between sham surgery and tMCAO for WT mice (e), and between WT and *Clec4e* tMCAO (f), show that, whether there are notable differences in microglial gene expression between sham surgery and tMCAO followed by 24 h reperfusion, *Clec4e* is not a contributing factor. Red dots: >1.5 upregulated in the first sample mentioned above graph (P<0.05); blue dots: >1.5 downregulated in the first sample mentioned above graph (P<0.05). (g) Absence of *Clec4e* mRNA in primary cortical neurons (left), cells of the brain endothelial cells line bEND.3 and primary astrocytes (right), untreated or under a time course of OGD (n=3). RNA from WT bone marrow-derived macrophages treated with LPS for 3 h is the positive control for all samples (n=1).

Close evaluation of Mincle expression, using both mRNA and protein analysis on WT and Clec4e^-/-^ mice, also failed to reveal Mincle expression by other cell types such as primary neurons, the brain endothelial cell line bEND.3, or primary astrocytes, including in response to OGD (Figure 5g). Mincle mRNA was also not detectable by qPCR in various human neuronal cell lines (CHP-212 or SH-SY5Y), nor under a variety of metabolic deprivation or inflammatory models (data not shown). We were unable to confirm the cellular pattern of Mincle protein expression in the mouse by immunohistochemistry (IHC), largely because of the poor specificity of anti-Mincle antibodies in mouse tissues (Supplementary Figure 1).

### In rat, Mincle is expressed on CD163-positive perivascular macrophages after permanent MCAO

In order to verify our observations that Mincle does not have widespread expression in the brain, we examined rat brain tissue after permanent MCAO. For this purpose, we carried out IHC on paraffin-embedded brain sections from rats that had undergone 24 h of permanent focal cerebral ischemia by distal occlusion of the middle cerebral artery^36^. Throughout the brain, in both infarct and non-ischemic tissue, Mincle^+^ cells were always associated with the vasculature. Monoclonal antibodies 4A9 and 16E3, generated in rat and mouse, respectively, provided the same result (Figure 6). Mincle was not present in cells positive for the pericyte marker alpha-SMA (Figure 6a,b,d), with the astrocyte marker GFAP (Figure 6a,c) or with the microglial marker Iba1 (Figure 6c). They always appeared sandwiched between pericytes and astrocytes (Figure 6b), and showed to be positive for CD163 (Figure 6d), which is considered a marker of rat brain perivascular macrophages^36^. We noted Mincle was readily visible in healthy and spontaneously hypertensive stroke-prone rats. Together the mouse and rat studies confirm that Mincle is not directly regulating microglia or astrocyte responses to sterile inflammation, but is restricted to macrophages associated with the brain vasculature. This pattern of expression may indicate a role for Mincle in regulating cerebral vasculature in response to injury, and thus help explain the apparent differences in phenotype of the Mincle KO mouse responding to brain MCAO or spinal cord neurotrauma.

**Figure 6.**
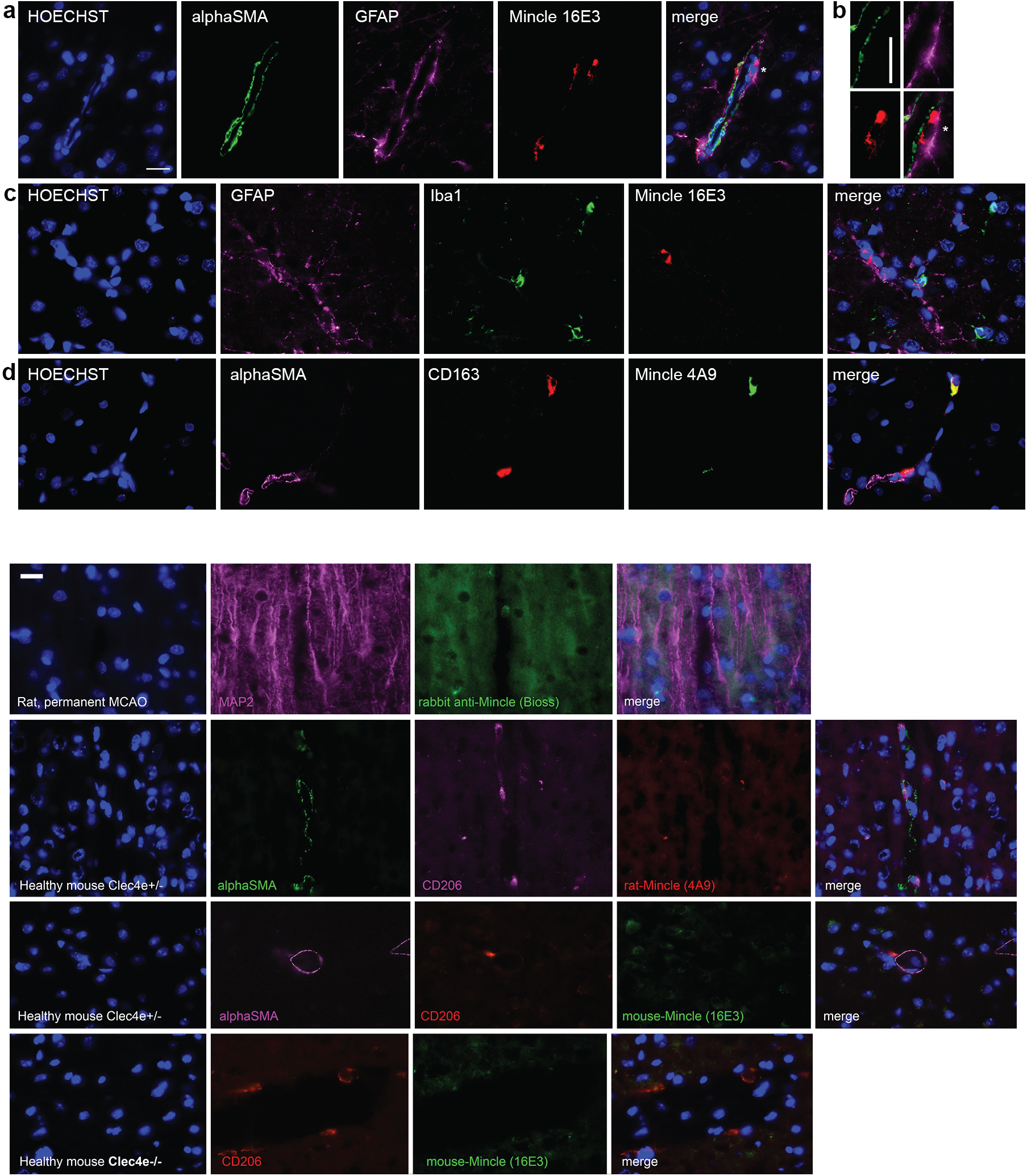
Mincle immunofluorescence in the cortical contralateral (non-infarcted) region of rats subjected to permanent MCAO shows presence of Mincle in the perivascular macrophages. (a) Mouse anti-Mincle antibody 16E3 shows Mincle^+^ cells do not co-localize with pericytes (alpha-SM^+^) or astrocytes (GFAP^+^), but with other cells located in the periphery of the blood vessels. (b) Detail of the previous cell (* indicates position) showing a Mincle^+^ cell (red) located between pericytes (green) and astrocytes (purple). (c) Mincle cells do not co-localize with microglia (Iba1^+^). (d) Rat anti-Mincle antibody 4A9 staining shows co-localisation with CD163, a marker of perivascular macrophages. Scale bars: 20 um.

## Discussion

The data presented here offer new insights into the initiation of sterile inflammation in the ischemic brain. Genetic deletion revealed a key role for Mincle in the severity of tissue damage following transient focal cerebral ischemia. This result can be mimicked with pharmacological inhibition of Syk, a downstream partner of Mincle, even when applied 3 h after the onset of reperfusion. Mincle’s major role is in the induction of ischemic inflammation, as the absence of Mincle in the brain was able to prevent the infiltration of peripheral immune cells, and to reduce the activation of neutrophils and monocytes present in the brain after transient focal cerebral ischemia in mice. Mincle was found in the brain perivascular macrophages of rat brain tissue, but in no other brain cell type. The resistance of the *Clec4e* mice to ischemic brain injury, but not to traumatic spinal cord injury, or peripheral (gut or heart) ischemia strongly suggests a unique role for Mincle in the context of cerebral ischemia.

Further clarification of Mincle’s role in ischemic stroke requires major improvements to the available molecular toolkit for mouse. We systematically evaluated the range of anti-Mincle antibodies, using our Mincle KO mouse to demonstrate specificity, and found serious deficits in their utility in mouse. These data are summarised in the supplementary files of this manuscript. It is on the strength of these antibodies that others have implicated Mincle in changes to innate immunity after ischemic stroke ^27^, subarachnoid haemorrhage^26^ and traumatic brain injury^39^. These studies are based on the presence of Mincle in neurons, and we have shown here by multiple molecular methods (RNA and protein, *in vitro* and *in vivo*) that Mincle was not present in mouse or rat neurons.

We also excluded a functional role for Mincle in microglia, which are radio-resistant and thus retained in the recipient animal in bone marrow chimeras^40^. When looking at TNF, a hallmark of the inflammatory phenotype, and in contrast with the dramatic differences between genotypes observed in tissue damage, we observed no differences in microglia-derived TNF production between WT and *Clec4e*^-/-^ animals after tMCAO. Microglia isolated from sham-operated or tMCAO brains one day after the injury did not express the *Clec4e* mRNA, and the transcriptional profile of Mincle-deficient microglia did not deviate from that of isogenic controls, after sham surgery or ischemia.

The studies that reported widespread Mincle expression in multiple brain cells also described a model for Mincle activation consistent with other peripheral immune receptors such as the TLR family^26,27^. That is, a model whereby Mincle acts as a necrotic cell receptor that is active on both CNS-resident cells and on peripheral immune cells recruited to the site of injury. It is possible that in these studies the reported increase in Mincle expression was due to influx of peripheral leukocytes. The concomitant upregulation of SAP130 in the ischemic hemisphere is puzzling because SAP130 is a constitutively expressed nuclear protein, which is not inducible like a cytokine, but instead is made available to immune receptors through cellular disruption via necrosis. Increased levels of SAP130 in whole tissue lysates are therefore most consistent with recruitment of more inflammatory cells to the lesion. The present data does not support the previously proposed model, directly disputing a model for Mincle as a peripherally-driven necrotic cell receptor. Firstly, the lack of Mincle-dependent phenotypes after spinal cord injury, and gut and heart ischemia, challenge the widely held model of Mincle as a peripheral myeloid receptor directing inflammatory responses to areas of necrotic cell damage. Secondly, the *Clec4e*^-/-^ mouse phenotype does not recapitulate that of other pattern recognition receptors with roles in necrotic cell recognition. For example, bone marrow chimera experiments using *Tlr2* or *Tlr4*, as well as double mutant mice showed that recipient KO mice were equally susceptible to tMCAO as WT mice. Functional benefit were only observed in WT donors receiving KO bone marrow, demonstrating that these receptors were most important in peripheral leukocyte activation^41^. In contrast, our own chimera studies demonstrated that the contribution of Mincle present in the peripheral immune cells had a relatively small impact on the severity of outcome after MCAO, which may be attributable to the relatively slow turnover of perivascular macrophages. On the other hand, loss of Mincle on CNS-resident was protective regardless of the genotype of the donor marrow. These data indicate that Mincle predominantly acts on an aspect specific to central nervous system architecture or function, rather than on the body’s ability to mount an inflammatory response to a sterile injury.

The significant reduction in brain infarct size in *Clec4e*^-/-^ animals suggests that blocking Mincle protects the cells in the penumbra region, that is, the brain region where blood perfusion is low but sufficient to maintain cell viability as a result of collateral blood vessel irrigation. Although we cannot fully exclude the possibility that there are gross anatomical differences in the development of the vasculature in *Clec4e*^-/-^ mice, the fact that blockade of Mincle signalling with a Syk inhibitor offered a similar functional benefit to genetic ablation of Mincle strongly argues against this and points towards functional differences in the response to the insult that are tissue-specific. Exhaustive analysis of spinal cord injury outcomes, in which the blood-CNS barrier is disrupted by mechanical forces at the outset, revealed no Mincle-dependant effect. This further supports our theory that Mincle acts via its influence over the microvasculature of the CNS, specifically in instances where its integrity is not directly compromised by the initiating event. Certainly, many features of the CNS response to an ischemic injury do not apply to SCI^42^.

The direct physical association between Mincle^+^ perivascular macrophages and the adventitial plane of alpha-SMA^+^ pericytes further supports a role for these cells in the regulation of the brain microvasculature and/or breakdown of the blood-brain barrier following an ischemic event. The dramatic injury reduction observed in *Clec4e*^-/-^ mice may therefore be explained by a key role for Mincle^+^ macrophages at this perivascular location with regards to injury propagation. An alternate and not mutually exclusive explanation would be that Mincle has a role in directing inflammatory cell recruitment across the blood-brain barrier, which may have contributed to the observed reduction in inflammatory infiltrate in the brains of *Clec4e*^-/-^ mice after stroke (Figure 2).

Although our results leave many unanswered questions about the role of Mincle in stroke, its dramatic influence over the outcome from brain ischemia and reperfusion injury, which mainly emerges from Mincle’s presence in the brain itself, warrants further investigation to resolve this aspect of the *Clec4e*^-/-^ phenotype for future therapeutic targeting.

## Acknowledgements

The authors wish to thank Professor Geoff R. Hill for contributions to the chimera experiments and valuable discussions on the study.

## Author contributions

CAW conception of project, experimental design and data generation (expression studies), sample provision, manuscript writing

TVA conception of project, experimental design, data generation, (MCAO and gut ischemia), manuscript writing

SM experimental design, data generation, manuscript writing

MJR data generation (spinal cord injury model), manuscript writing

MG experimental design, data generation (cytokine data in ischemic mice)

KPAM data generation (bone marrow chimera studies)

MF data generation (cardiac ischemia), data interpretation

ES data generation (cardiac ischemia)

OK data analysis (microarray)

DD Sample provision, data interpretation

IMM Sample provision, data interpretation

RBA Sample provision

SCT Sample provision

YLL Sample provision

PJB data generation (spinal cord injury model)

NAR data generation (cardiac ischemia)

TM experimental design (Cytokine data in ischemic mice)

YHH (deceased) data generation (molecular characterisation of Mincle)

**Supplementary Figure 1.** Mincle antibody staining on rat and mouse brain. (a) The rabbit anti-Mincle antibody (Bioss) did not reveal any clear signal on rat brain tissue. (b,c) Using alpha-SMA as a marker of pericytes and CD206 as a marker of perivascular macrophages, both rat anti-Mincle and mouse anti-Mincle antibodies reveal no specific staining in healthy brains from *Clec4e* mice. (d) Background staining could be appreciated also in *Clec4e* brains. Scale bar: 20 um.

**Supplementary Figure 2.** The ARRIVE guidelines checklist for this study.

|  | ITEM | RECOMMENDATION | Section/<br>Paragraph |
| --- | --- | --- | --- |
| Title | 1 | Provide as accurate and concise a description of the content of the article as possible. | Title |
| Abstract | 2 | Provide an accurate summary of the background, research objectives, including details of the species or strain of animal used, key methods, principal findings and conclusions of the study. | Abstract |
| **INTRODUCTION** |  |  |  |
| Background | 3 | a. Include sufficient scientific background (including relevant references to previous work) to understand the motivation and context for the study, and explain the experimental approach and rationale.<br>b. Explain how and why the animal species and model being used can address the scientific objectives and, where appropriate, the study's relevance to human biology. | a) Paragraph 1-3<br>b) Paragraph 3-4 |
| Objectives | 4 | Clearly describe the primary and any secondary objectives of the study, or specific hypotheses being tested. | Paragraph 4 |
| **METHODS** |  |  |  |
| Ethical statement | 5 | Indicate the nature of the ethical review permissions, relevant licences (e.g. Animal [Scientific Procedures] Act 1986), and national or institutional guidelines for the care and use of animals, that cover the research. | Paragraph 1 |
| Study design | 6 | For each experiment, give brief details of the study design including:<br>a. The number of experimental and control groups.<br>b. Any steps taken to minimise the effects of subjective bias when allocating animals to treatment (e.g. randomisation procedure) and when assessing results (e.g. if done, describe who was blinded and when).<br>c. The experimental unit (e.g. a single animal, group or cage of animals).<br>A time-line diagram or flow chart can be useful to illustrate how complex study designs were carried out. | a,b)<br>Paragraph 1, 3, 6, 7, 9, 10<br>c) Paragraph 15 |
| Experimental procedures | 7 | For each experiment and each experimental group, including controls, provide precise details of all procedures carried out. For example:<br>a. How (e.g. drug formulation and dose, site and route of administration, anaesthesia and analgesia used [including monitoring], surgical procedure, method of euthanasia). Provide details of any specialist equipment used, including supplier(s).<br>b. When (e.g. time of day).<br>c. Where (e.g. home cage, laboratory, water maze).<br>d. Why (e.g. rationale for choice of specific anaesthetic, route of administration, drug dose used). | a,d)<br>Paragraph 1, 3, 6, 7, 9, 10<br>b)<br>Paragraph 1 |
| Experimental animals | 8 | a. Provide details of the animals used, including species, strain, sex, developmental stage (e.g. mean or median age plus age range) and weight (e.g. mean or median weight plus weight range).<br>b. Provide further relevant information such as the source of animals, international strain nomenclature, genetic modification status (e.g. knock-out or transgenic), genotype, health/immune status, drug or test naïve, previous procedures, etc. | a) Paragraph 1, 3, 6, 7, 9, 10<br>b) Paragraph 1 |
| Housing and husbandry | 9 | Provide details of:<br>a. Housing (type of facility e.g. specific pathogen free [SPF]; type of cage or housing; bedding material; number of cage companions; tank shape and material etc. for fish).<br>b. Husbandry conditions (e.g. breeding programme, light/dark cycle, temperature, quality of water etc for fish, type of food, access to food and water, environmental enrichment).<br>c. Welfare-related assessments and interventions that were carried out prior to, during, or after the experiment. | a,b)Paragraph 1<br>c)Paragraph 1, 3, 6, 7, 9, 10 |
| Sample size | 10 | a. Specify the total number of animals used in each experiment, and the number of animals in each experimental group.<br>b. Explain how the number of animals was arrived at. Provide details of any sample size calculation used.<br>c. Indicate the number of independent replications of each experiment, if relevant. | a) Figures 1-5 (individual values plotted), Fig 1 legend<br>b)Paragraph 15 |
| Allocating animals to experimental groups | 11 | a. Give full details of how animals were allocated to experimental groups, including randomisation or matching if done.<br>b. Describe the order in which the animals in the different experimental groups were treated and assessed. | a,b)Paragraph 1, 3, 6, 7, 9, 10 |
| Experimental outcomes | 12 | Clearly define the primary and secondary experimental outcomes assessed (e.g. cell death, molecular markers, behavioural changes). | a,b) Paragraph 1, 3, 6, 7, 9, 10 |
| Statistical methods | 13 | a. Provide details of the statistical methods used for each analysis.<br>b. Specify the unit of analysis for each dataset (e.g. single animal, group of animals, single neuron).<br>c. Describe any methods used to assess whether the data met the assumptions of the statistical approach. | Paragraph 15 |
| **RESULTS** |  |  |  |
| Baseline data | 14 | For each experimental group, report relevant characteristics and health status of animals (e.g. weight, microbiological status, and drug or test naïve) prior to treatment or testing. (This information can often be tabulated). | Methods paragraph 1 |
| Numbers analysed | 15 | a. Report the number of animals in each group included in each analysis. Report absolute numbers (e.g. 10/20, not 50%<sup>2</sup>).<br>b. If any animals or data were not included in the analysis, explain why. | Methods paragraph 2 |
| Outcomes and estimation | 16 | Report the results for each analysis carried out, with a measure of precision (e.g. standard error or confidence interval). | Figures 1-5 |
| Adverse events | 17 | a. Give details of all important adverse events in each experimental group.<br>b. Describe any modifications to the experimental protocols made to reduce adverse events. | n.a. |
| **DISCUSSION** |  |  |  |
| Interpretation/scientific implications | 18 | a. Interpret the results, taking into account the study objectives and hypotheses, current theory and other relevant studies in the literature.<br>b. Comment on the study limitations including any potential sources of bias, any limitations of the animal model, and the imprecision associated with the results<sup>2</sup>.<br>c. Describe any implications of your experimental methods or findings for the replacement, refinement or reduction (the 3Rs) of the use of animals in research. | Throughout discussion |
| Generalisability/translation | 19 | Comment on whether, and how, the findings of this study are likely to translate to other species or systems, including any relevance to human biology. | Throughout discussion |
| Funding | 20 | List all funding sources (including grant number) and the role of the funder(s) in the study. | Detailed in “sources of support” |

## References

1. Woodruff, T. M. et al. Pathophysiology, treatment, and animal and cellular models of human ischemic stroke. Mol Neurodegener 6, 11 (2011).

2. Chamorro, A. et al. The immunology of acute stroke. Nat Rev Neurol 8, 401–410 (2012).

3. Murray, K. N., Buggey, H. F., Denes, A. & Allan, S. M. Systemic immune activation shapes stroke outcome. Mol. Cell. Neurosci. 53, 14–25 (2013).

4. Tang, S. C. et al. Pivotal role for neuronal Toll-like receptors in ischemic brain injury and functional deficits. Proc Natl Acad Sci U S A 104, 13798–13803 (2007).

5. Heiman, A., Pallottie, A., Heary, R. F. & Elkabes, S. Toll-like receptors in central nervous system injury and disease: A focus on the spinal cord. Brain, Behavior, and Immunity 42, 232–245 (2014).

6. Downes, C. E. et al. MyD88 Is a Critical Regulator of Hematopoietic Cell-Mediated Neuroprotection Seen after Stroke. PLoS One 8, e57948 (2013).

7. Vartanian, K. B. et al. LPS preconditioning redirects TLR signaling following stroke: TRIF-IRF3 plays a seminal role in mediating tolerance to ischemic injury. J Neuroinflammation 8, 140 (2011).

8. Kim, D. et al. A critical role of toll-like receptor 2 in nerve injury-induced spinal cord glial cell activation and pain hypersensitivity. J. Biol. Chem. 282, 14975– 14983 (2007).

9. Kigerl, K. A. et al. Toll-like receptor (TLR)-2 and TLR-4 regulate inflammation, gliosis, and myelin sparing after spinal cord injury. J. Neurochem. 102, 37–50 (2007).

10. Brea, D. et al. Toll-like receptors 2 and 4 in ischemic stroke: outcome and therapeutic values. J Cereb Blood Flow Metab 31, 1424–1431 (2011).

11. Maestrini, I. et al. Higher neutrophil counts before thrombolysis for cerebral ischemia predict worse outcomes. Neurol. 85, 1408–1416 (2015).

12. Holloway, P. M. et al. Both MC1 and MC3 Receptors Provide Protection From Cerebral Ischemia-Reperfusion–Induced Neutrophil Recruitment. Arterioscler. Thromb. Vasc. Biol. 35, 1936–1944 (2015).

13. Mori, E., del Zoppo, G. J., Chambers, J. D., Copeland, B. R. & Arfors, K. E. Inhibition of polymorphonuclear leukocyte adherence suppresses no-reflow after focal cerebral ischemia in baboons. Stroke 23, 712–718 (1992).

14. Neumann, J. et al. Very-late-antigen-4 (VLA-4)-mediated brain invasion by neutrophils leads to interactions with microglia, increased ischemic injury and impaired behavior in experimental stroke. Acta Neuropathol. 129, 259–277 (2015).

15. Lalancette-Hébert, M., Gowing, G., Simard, A., Weng, Y. C. & Kriz, J. Selective ablation of proliferating microglial cells exacerbates ischemic injury in the brain. J. Neurosci. 27, 2596–605 (2007).

16. Russo, M. V & McGavern, D. B. Immune Surveillance of the CNS following Infection and Injury. Trends Immunol. 36, 637–50 (2015).

17. Gensel, J. C. et al. Toll-Like Receptors and Dectin-1, a C-Type Lectin Receptor, Trigger Divergent Functions in CNS Macrophages. J. Neurosci. 35, 9966–76 (2015).

18. Yamasaki, S. et al. Mincle is an ITAM-coupled activating receptor that senses damaged cells. Nat Immunol 9, 1179–1188 (2008).

19. Matsumoto, M. et al. A novel LPS-inducible C-type lectin is a transcriptional target of NF-IL6 in macrophages. J. Immunol. 163, 5039–5048 (1999).

20. Schoenen, H. et al. Cutting edge: mincle is essential for recognition and adjuvanticity of the mycobacterial cord factor and its synthetic analog trehalose-dibehenate. J Immunol 184, 2756–2760 (2010).

21. Vijayan, D., Radford, K. J., Beckhouse, A. G., Ashman, R. B. & Wells, C. A. Mincle polarizes human monocyte and neutrophil responses to Candida albicans. Immunol Cell Biol 90, 889–895 (2012).

22. Wells, C. A. et al. The macrophage-inducible C-type lectin, mincle, is an essential component of the innate immune response to Candida albicans. J Immunol 180, 7404–7413 (2008).

23. Drummond, R. A., Saijo, S., Iwakura, Y. & Brown, G. D. The role of Syk/CARD9 coupled C-type lectins in antifungal immunity. Eur J Immunol 41, 276–281 (2011).

24. Strasser, D. et al. Syk kinase-coupled C-type lectin receptors engage protein kinase C-sigma to elicit Card9 adaptor-mediated innate immunity. Immunity 36, 32–42 (2012).

25. Andre, P. et al. Critical role for Syk in responses to vascular injury. Blood 118, 5000–5010 (2011).

26. He, Y. et al. Macrophage-Inducible C-Type Lectin/Spleen Tyrosine Kinase Signaling Pathway Contributes to Neuroinflammation After Subarachnoid Hemorrhage in Rats. Stroke 46, 2277–2286 (2015).

27. Suzuki, Y. et al. Involvement of Mincle and Syk in the changes to innate immunity after ischemic stroke. Sci. Rep. 3, 3177 (2013).

28. Ishikawa, E. et al. Direct recognition of the mycobacterial glycolipid, trehalose dimycolate, by C-type lectin Mincle. J Exp Med 206, 2879–2888 (2009).

29. Bederson, J. B. et al. Rat middle cerebral artery occlusion: evaluation of the model and development of a neurologic examination. Stroke. 17, 472–476 (1986).

30. Gelderblom, M. et al. Temporal and spatial dynamics of cerebral immune cell accumulation in stroke. Stroke 40, 1849–1857 (2009).

31. Harrison, M. et al. Vertebral landmarks for the identification of spinal cord segments in the mouse. Neuroimage 68, 22–29 (2013).

32. Blomster, L. V. et al. Mobilisation of the splenic monocyte reservoir and peripheral CX3CR1 deficiency adversely affects recovery from spinal cord injury. Exp. Neurol. 247, 226–240 (2013).

33. Brennan, F. H. et al. The Complement Receptor C5aR Controls Acute Inflammation and Astrogliosis following Spinal Cord Injury. J. Neurosci. 35, 6517–31 (2015).

34. Basso, D. M. et al. Basso Mouse Scale for locomotion detects differences in recovery after spinal cord injury in five common mouse strains. J. Neurotrauma 23, 635–659 (2006).

35. Pamuk, O. N. et al. Spleen tyrosine kinase inhibition prevents tissue damage after ischemia-reperfusion. Am. J. Physiol. Gastrointest. Liver Physiol. 299, G391–399 (2010).

36. Tarr, D. et al. Hyperglycemia accelerates apparent diffusion coefficient-defined lesion growth after focal cerebral ischemia in rats with and without features of metabolic syndrome. J. Cereb. Blood Flow Metab. 33, 1556–1563 (2013).

37. Wells, C. A. et al. Stemformatics: Visualisation and sharing of stem cell gene expression. Stem Cell Res. 10, 387–395 (2012).

38. Smyth, G. K. in Bioinformatics and Computational Biology Solutions using R and Bioconductor 397–420 (2005). doi:10.1007/0-387-29362-0_23

39. de Rivero Vaccari, J. C. et al. Mincle Signaling in the Innate Immune Response after Traumatic Brain Injury. J. Neurotrauma 236, 228–236 (2014).

40. Gomez Perdiguero, E., Schulz, C. & Geissmann, F. Development and homeostasis of ‘resident’ myeloid cells: the case of the microglia. Glia 61, 112– 120 (2013).

41. Shichita, T. et al. Peroxiredoxin family proteins are key initiators of post-ischemic inflammation in the brain. Nat Med 18, 911–917 (2012).

42. Iadecola, C. & Anrather, J. Stroke research at a crossroad: asking the brain for directions. Nat. Neurosci. 14, 1363–1368 (2011).

